# Plant Accessible Tissue Clearing Solvent System (PATCSOS) for 3-D Imaging of Whole Plants

**DOI:** 10.1101/2024.04.20.590386

**Authors:** Hantao Zhang, Lei Zhu, Hu Zhao, Zhen Li

## Abstract

Tissue clearing is a technique to make the inner structure of opaque tissue visible to achieve 3-dimensional (3-D) tissue imaging by unifying the refractive indexes of most of the cell components. Tissue clearing is widely used in animal tissue imaging, where whole body 3-D imaging has been realized. However, it has not been widely used in plant research. Most plant tissue clearing protocols have their disadvantages, including low efficiency, not being fluorescence-friendly and poor transparency on tissues with a high degree of lignification. In this work, we developed a new plant tissue clearing method for whole plant imaging, named Plant Accessible Tissue Clearing Solvent System (PATCSOS), which was based on the Polyethylene Glycol-associated Solvent System (PEGASOS). The PATCSOS method realized extensive transparency of plant tissues, including the flower, leaf, stem, root, and seed of Arabidopsis thaliana, with high efficiency. The PATCSOS method consists of four main steps: fixation, decolorization/delipidation, dehydration, and clearing. Subsequently a rapid and efficient clearing of mature plant tissue can be achieved. With PATCSOS, we can image Arabidopsis seedling in their entirety in 3-D using endogenous cellulose autofluorescence. What’s more, the PATCSOS method is compatible with fluorescence protein imaging and GUS staining, which greatly expands the applicability of this method. We also imaged intact *Nicotiana benthamiana* leaf and *Zea mays* embryos. Our results showed that the PATCSOS clearing method is an excellent tool to study plant development and cell biology.

## 1 Introduction

To better investigate plant growth and development, it is important to obtain 3-D images of plant tissues and observe the inner structure of the plants. Different cell components have different refractive indices (RI), for example, water has a RI of 1.33, lipids have a RI of above 1.45, proteins have a RI of above 1.44, plant cell walls have a RI of 1.42, and oxygen produced by green plants has a RI of about 1.00[1-5]. As a result of these differences, the plant tissues appear to us to be opaque.

To address the issue of refractive index differences, the tissue transparency technique was adopted a method of making the entire tissue transparent by unifying the RI of most cellular components. Existing tissue clearing methods can be divided into two categories: organic solvents and aqueous solution system[5, 6]. For example, Visikol and Benzyl benzoate/benzyl alcohol (BABB) belong to the organic solvent system [5, 7, 8]. In the BABB system, organic solvents, including benzyl alcohol and benzyl benzoate, were used. iTOMEI, ClearSee and its derivative clearing methods, and Pea-CLARITY used aqueous solutions[9]. These methods have many disadvantages, such as poor transparency, long processing times, not being fluorescence-friendly, and, most importantly, not being optimized for clearing plant tissues.

In 2018, Hu Zhao and his team developed an effective tissue clearing method for mouse whole body imaging called PEGASOS. PEGASOS can clear a wide range of mouse tissues and PEGASOS system can protect the fluorescence signal from being quenched [4]. Based on PEGASOS, we developed a plant accessible tissue clearing solvent system (PATCSOS) and achieved considerable transparent effect for the whole plant of *Arabidopsis*, which can warrant the construction of 3-D images of plant tissues using autofluorescence from cellulose. We further confirmed that the modified PEGASOS method is compatible with imaging with fluorescence protein and GUS staining in plants.

## 2 Materials and Methods

### 2.1 Plant materials and growth condition

*Arabidopsis thaliana* (*A. thaliana*) accession *Columbia-0* (*Col-0*) and transgenic line *ANTPro::GFP, DR5::GUS* [10] and *HB29Pro::GUS* were constructed (see below)The *Arabidopsis* seeds were surface-sterilized for 15 min in 0.5% sodium hypochlorite and sown on 1/2 Murashige & Skoog medium containing 0.9% plant TC agar (PhytoTechnology Laboratories, Shawnee Mission, KS, USA). The *Arabidopsis* grew at 22°C under a 16 h: 8 h, light: dark photoperiod.

### 2.2 Molecular cloning and transformation

A 2-kb region of the *HB29* (AT1G69600) promoter and 2-kb region of the *ANT* (AT4G37750) promoter were amplified. *HB29* promoter was inserted into pCAMBIA1391. *ANT* promoter, *GFP* gene and *NOS* terminator were inserted into pCAMBIA1300 vectors to prepare *HB29 Pro::GUS* and *ANT Pro::GFP* constructs. *HB29 Pro::GUS* and *ANT Pro::GFP* constructs were generated and transformed into *Col-0* plants to produce the marker lines. The plant transformation vector pCAMBIA1300 and pCAMBIA1391 was used to generate transformed plants. Plants were transformed using Agrobacterium tumefaciens strain GV3101 using the floral dip method. [11]. The sequences of primers were listed in Table S1.

### 2.3 Preparation of PATCSOS solutions

The clearing solutions was adopted from PEGASOS with some modifications[4].

*Fixation solution*. Fixation solution was prepared by mixing 4% (w/v) Paraformaldehyde (PFA, Sigma-Aldrich 158127) with 0.01 M PBS (pH 7.4, Solarbio P1003).

*Gradient tB solution*.30%, 50% and 70% Tert-Butanol (tB, Aladdin T119717) in water v/v and 3% w/v Quadrol (Sigma-Aldrich 122262) was added afterwards.

*tB-PEG dehydration solution*. 70% v/v Tert-Butanol (Aladdin T119717), 27% v/v PEG methacrylate Mn 500 (PEGMMA500, Sigma-Aldrich 447943) and 3% w/v Quadrol (Sigma-Aldrich 122262).

*BB-PEG clearing medium*. 75% v/v benzyl benzoate (BB, Aladdin B400547), 25% v/v PEGMMA500 (Sigma-Aldrich 447943), and 3% w/v Quadrol (Sigma-Aldrich 122262).

### 2.4 Passive immersion procedure

#### Seedlings

*Arabidopsis* seedlings were immersed in 1 mL fixation solution and fixed overnight at room temperature. The seedlings were washed with 1 mL of ddH_2_O at 37°C under gentle shaking for 20 minutes and repeated three times. Then they were immersed in a 30% tB solution for one day at 37°C under gentle shaking to remove the chlorophyll. Afterwards, the 30% tB solution was replaced with 50% tB for one day and then with 70% tB for another day, both at 37°C under gentle shaking. The tissues were placed in a dehydration solution overnight at 37°C under gentle shaking. Finally, the dehydrated tissues were immersed in clearing medium for at least 24 h at 37°C under gentle shaking.

#### Mature tissues

For mature tissues, the procedures were modified to fit the large diameter and cellulose rich nature of the tissue. For bolting *Arabidopsis*, the inflorescence stem was immersed in 4% paraformaldehyde (PFA) overnight at room temperature, the tissues were washed three times with ddH_2_O, and then immersed in 30% tB for two days, and the 30% tB solution was replaced twice a day to better remove chlorophyll. The stems were immersed in 50% tB for one day, 70% tB for one day, and a dehydration solution for one day. And at last, they were immersed in the clearing medium for at least one day. All above, the steps were performed on a 37°Cshaker.

### 2.5 GUS Staining

Seven-day-old *HB29pro::GUS* seedlings were used to test compatibility between PATCSOS and GUS staining. Seedlings were immersed in GUS assay solution (Table S2) at 37°C in the dark for 4 hours using a widely adapted protocol [12].

In the decolorizing step, the seedlings were immersed in 30% tB for 10 minutes, 50% tB for 10 minutes and 70% tB for 10 minutes to adapt to the PATCSOS system, while in traditional methods, ethanol and acetate solution was used.

### 2.6 Microscopy and image analysis

GUS imaging was acquired on a Leica AF 6000 stereoscopic microscopy.

3-D fluorescence imaging of plant tissues was acquired on a Leica STELLARIS 5 confocal microscopy (Laser lines: 488, 561, 638; Leica HyD photon detector).

Raw images were collected in lossless 8-bit TIFF format. Single GFP fluorescence images were processed with Fiji (NIH). 3-D reconstructed images were generated using Imaris 9.0 (Bitplane). Stack images were generated using the “volume rendering” function. 3-D image clipping was realized using the “Clipping Plane” function. Background blocking was realized using the “Oblique Slicer” and “Ortho Slicer” function. 3-D pictures were generated using the “Snapshot” function. Movies were made using the “Animation” function.

## 3 Results

### 3.1 Optimization of the PATCSOS procedure and evaluation of clearing efficiency on plant tissues

There are five to six steps in the PEGASOS protocol, i.e.: fixation, (decalcification), decolorization, delipidation, dehydration, and clearing [4]. The decalcification step was omitted due to the absence of bone in the plant tissue. The decolorization step can also be omitted as there is no heme in the plant tissue and the gradient tB solution also has the effect of removing chlorophyll, therefore the steps of delipidation and decolorization can be combined to reduce the complexity of the method. The optimized PATGSOS protocol for plant tissue clearing included the following steps: fixation, decolorization/delipidation, dehydration, and clearing.

Mature *Arabidopsis* inflorescence stems (Fig. 1b, 1e) became completely transparent after 7 days in the clearing solution, the clearing duration is even shorter for seedlings (Fig. 1a, 1d). After clearing, the inner vascular bundles can be seen clearly in the inflorescence stem, and seeds in silique are also visible. This shows that the pericarp has been completely cleared. Thinner plant tissues, such as leaves and flowers, got the best transparent effect; they were completely invisible in the clearing solution after processing (Fig. 1e).

**Figure 1.**
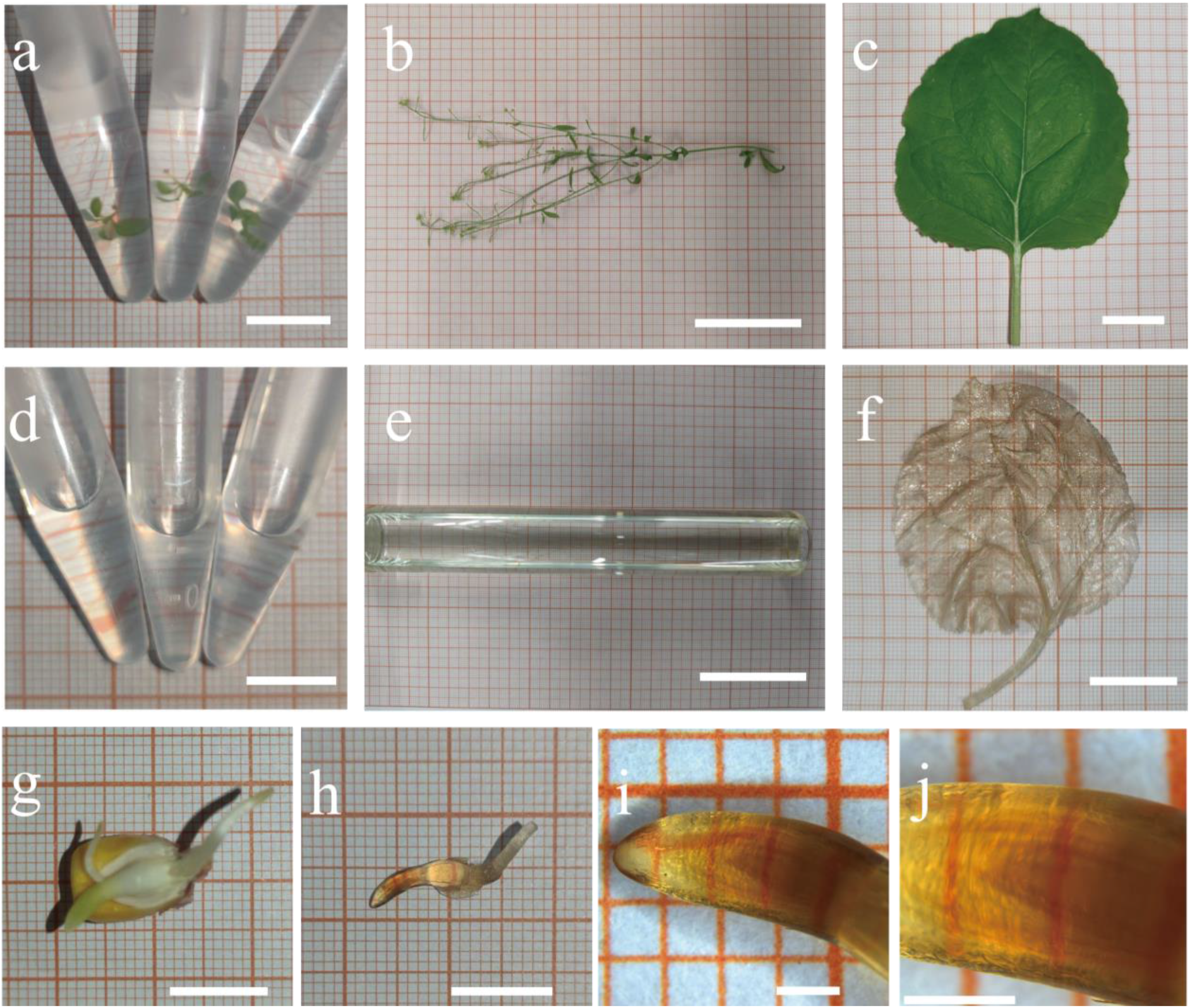
Different plant materials treated by PATCSOS. *Arabidopsis* seedlings were visible in PFA (a) without clearing, and after PATCSOS treatment, the seedlings became invisible (d). So as the mature *Arabidopsis* plant, a part of the inflorescence stem(b) also can be cleared by PATCSOS (e). *Nicotiana tabacum* leaf (c) and germinated *Zea mays* embryo (g) before (c, g) and after (f, h) processed with PEGASOSFP. At the center of the embryo, the developing plumule can be seen (i, j). Bar, 1 cm (a, d, g, h); bar, 5 cm (b, e); bar, 2 cm (c, f); bar, 1 mm (i, j).

We also cleared *Nicotiana tabacum* leaf and *Zea mays* embryo after germination. The chlorophyll in tobacco leaf was removed effectively, and the whole leaf was totally transparent after clearing (Fig. 1c, 1f). The embryo and endosperm of corn seed was removed after clearing to get a better view of the corn embryo (Fig. 1g,1 h). And after magnifying the embryo with a stereoscopic microscope, we can see the developing plumule inside the coleoptile (Fig. 1i, 1j).

### 3.2 3-D imaging with autofluorescence

Autofluorescence is a normal phenomenon in plant tissues. Many cell components such as chlorophyll and cellulose can emit autofluorescence. Autofluorescence is generally regarded as interference signal in fluoresce imaging and various methods have been developed to remove it [13, 14]. But it serves as a natural marker to show the cell profile and plant tissue structure. After clearing, we can capture autofluorescence signal from cellulose clearly, with which a 3-D image of plant tissue was obtained (Fig. 2).

**Figure 2.**
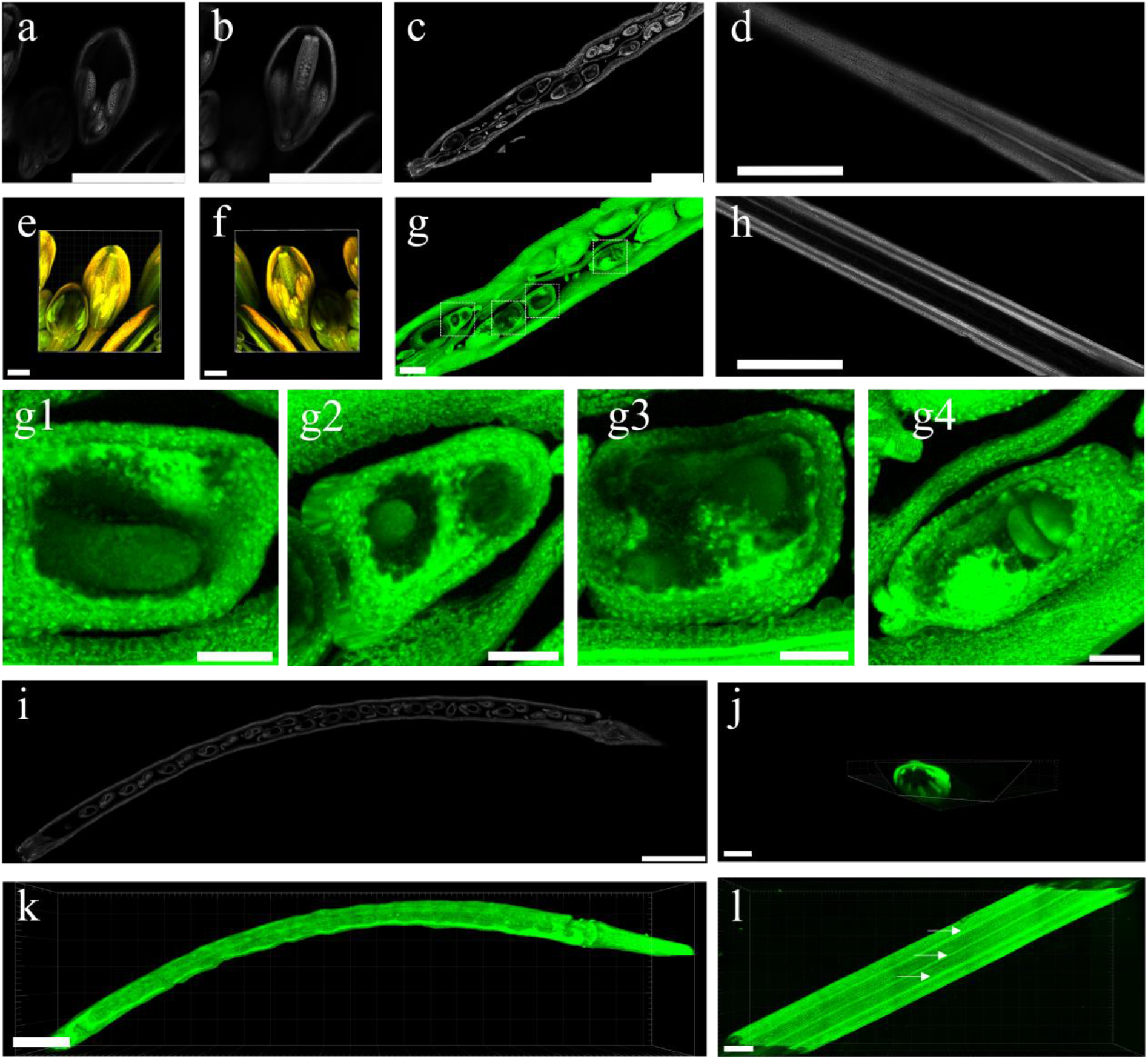
3-D imaging of *Arabidopsis* tissues using autofluorescence: (a-b) two images of single light slices of flowers with different focal planes; (e-f) 3-D reconstructed images of flowers; (c) single slice of silique; (g) 3-D reconstructed image to highlight the embryo; g1-g4: magnification of the image framed by the white dotted line in (g); (i) single slice of a mature silique; (k) 3-D reconstructed image of the mature silique in (i); (d, h) single slices of inflorescence stem focused on different depth of focus; (j, l) different view of the inflorescence stem 3-D image, and vascular bundles are labeled by arrows in (l). Most autofluorescence was excited using 488-nm argon lasers, and the flower tissue imaging was also excited using 561-nm laser. Bar, 1 mm (a, b, c, d, h, i, k); bar, 300 μm (e, f, g, l); bar, 100 μm (g1, g2, g3, g4); bar, 500 μm (j).

We imaged flower of *Arabidopsis* with autofluorescence using 10×objective in two channels (excitation: 488 nm and 561 nm), the step size was set at 1 μm. Two single slices at different Z-axis positions showing pollens in the anther sac and ovules in the ovary, respectively (Fig. 2a, 2b). Clear observation of such tissues indicated applicability of the PATCSOS for clearing flowers tissues. Reconstructed 3-D images of flowers (Fig. 2e, 2f) can help us track the direction of the vascular system from pedicel to stamens and pistil of the flower. So, the PATCSOS system can realize high resolution 3-D imaging of fine structures of plants, regardless of inner or outer region without dissection. Whole tissue clearing and 3-D imaging is more intuitive and can maintain the relative position of each part compared with traditional dissection methods [15].

More tender and more mature silique tissues were also imaged. Fig. 2c is one of the slices of the tender silique captured with 40×objective stimulated by 488 nm laser with a step size of 1μm. Because of limited working distance of objective, we only imaged half of a silique. The slicer tool in Imaris was used to reveal inner structure of the silique, and the development condition of the embryo can be clearly observed through the reconstructed image (Fig. 2g). Seeds in different developmental stage and their relative positions in the silique can be clearly visualized (Fig. 2g1-g4).

A 20× objective with long working distance was used to image another more mature silique with a small piece of stipe (Fig. 2i). Regretfully, because of the flavonoid of the testa, the inner structure of some seeds was not clearly visible (Fig. 2i). Images and movie of silique in 3-D view showed continuous change from the outer silique to the inner, and we can see the arrangement of seeds in a silique (Fig 2k, Movie S1).

Part of inflorescence stem was imaged with a 10×objective stimulated by 488 nm laser, the step size was set as 1 μm (Fig. 2d, 2h, 2j, 2l). The structure of vascular bundles was clearly countable from the rendered 3-D image (Fig. 2l). The diameter of the inflorescence stem was about 1 mm, the tissue needs to be highly transparent to be imaged at such high resolution and quality. This result showed that the PATCSOS protocol has good clearing effect on *Arabidopsis* tissues imaging.

### 3.3 GFP fluorescence imaging

Fluorescence protein (FP) labelling is widely used to investigate tissue development and protein subcellular localization in cell biology [5]. In whole tissue clearing analysis, the ultimate goal was to image the inner structure of tissues using fluorescence;thus, the protection of fluorescent proteins during clearing is of critical importance.

The passive immerse process employed by PATCSOS can readily protect the fluorescent proteins by using gradient tB to replace ethanol in traditional plant material processing methods and adding Quadrol to keep a basic environment to maintain activities of the FPs. However, GFP imaging of *Arabidopsis* line *ANT Pro::GFP* presented the dilemma of discriminating fluorescence signal between autofluorescence and GFP. Fluorescence from FPs can only be stimulated by excitation light with specific wavelength. In plants, chlorophyll and lignin stand out as the key auto fluorescent substances, yet a variety of additional compounds exhibit autofluorescence when subjected to ultraviolet or visible light excitation, encompassing elements from the cytoplasm and the structural cell walls [16]. While for autofluorescence, a broad wavelength range can be used for excitation. Thus, during image acquisition, we collected data from two channels, the first channel used 488 nm laser for excitation and signals from both GFPs and autofluorescence were acquired, the other channel used 561 nm laser for excitation and signal only from autofluorescence was acquired. Images from these two channels were subtracted to obtain images only from the fluorescence proteins [17].

Fluorescence images from one section of *Arabidopsis ANTPro::GFP* stipe was imaged with excitation laser at 488 and 561 nm, the images showed distinct distribution of fluorescence signals from GFP and autofluorescence (Fig. 3a, 3b). When a 488 nm laser was used, fluorescence from both GFP and autofluorescence can be clearly visualized from cambium and epidermis region of the stipe (Fig. 3a). When a 561 nm laser was used, fluorescence only from epidermis can be imaged (Fig. 3b). Since *ANT* is a marker gene to label the cambium[18], the *ANTPro::GFP* construct should emit fluorescence signals only from the cambium region, but fluorescence image from the 488 nm channel showed ubiquitous distribution of signal due to interference from autofluorescence from cellulose (Fig. 3a). We further merged images from the two channels (Fig. 3d), to distinguish autofluorescence from GFP signal, the orange region represents autofluorescence signal from the 488 nm and 561 nm channels and the green region represents GFP signal from only the 488 nm channel.

**Figure 3.**
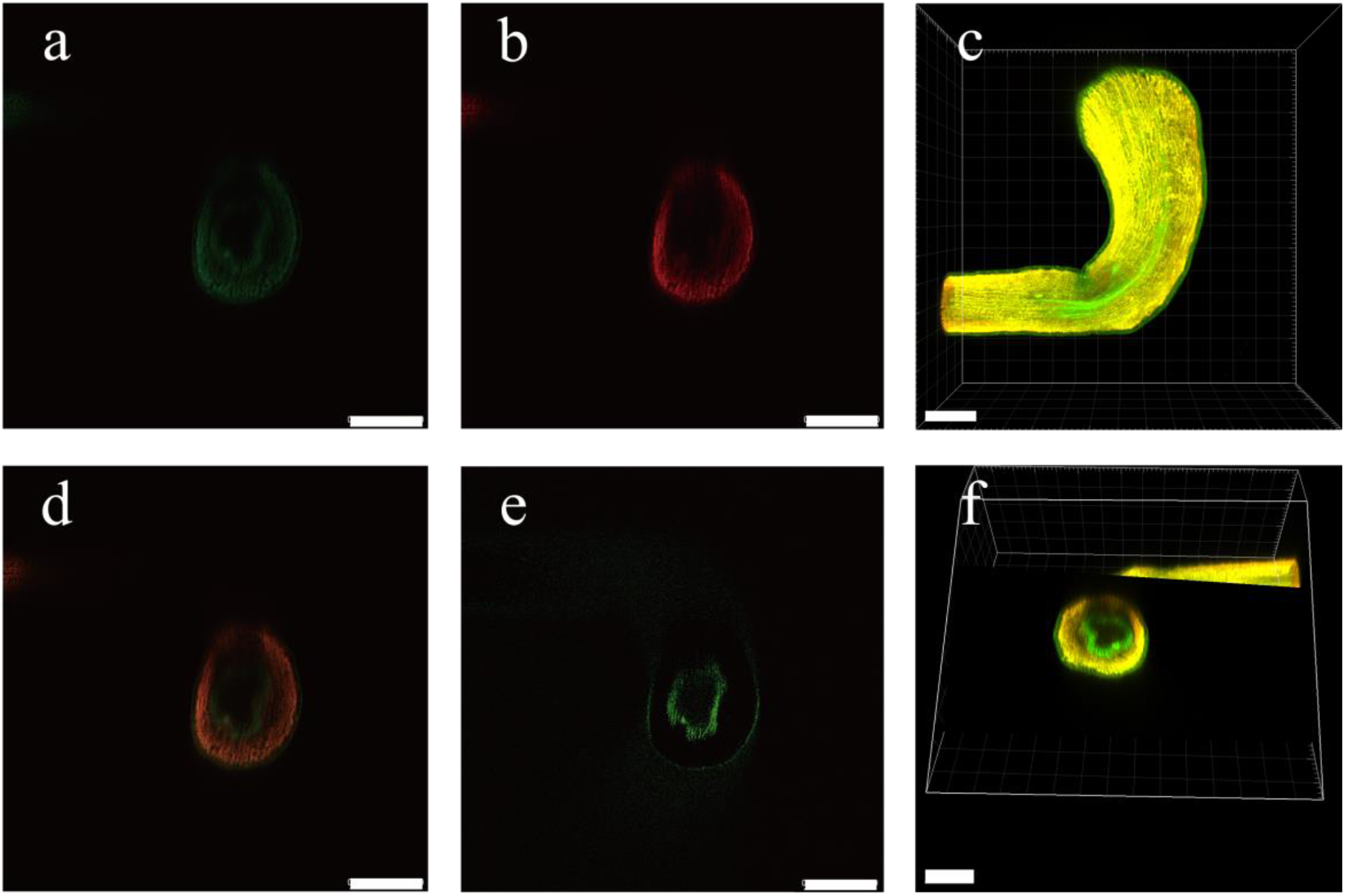
Confocal imaging of the stipe of *ANTPro::GFP* transgenic *Arabidopsis*. The GFP fluorescence (a) and the autofluorescence (b) imagines are captured. And merge (d), subtract (e) and view them in 3-D vision (c, f). Two channels were used for imaging. Channel 1 used 488-nm laser as exciting light, and emission light was collected from 499.8-579.3 nm. Channel 2 used 561-nm laser as exciting light, and emission light was collected from 623.5-725.7 nm. Bar, 250 μm (a, b, d, e); bar, 200 μm (c, f).

We further used “Subtract” function of the Fiji software remove autofluorescence interference signal. The brightness of the two images was adjusted to the same level, and the “Subtract” function was used to performance subtraction between Fig. 3a and 3b, which resulted in a clean GFP fluorescence image of *Arabidopsis* cambium (Fig. 3e).

Through 3-D imaging reconstruction and background subtraction, a constant *ANT* express pattern in the whole stipe was observed (Fig. 3c, 3f), where, the yellow part represents autofluorescence and the green part represents GFP signal, and the cambium pattern in the stipe can be clearly visualized.

### 3.4 Whole seedling 3-D imaging

3-D imaging of whole *Arabidopsis* seedling was also realized with PATCSOS. The immature development status of cell wall makes the seedling tissue tenderer than mature inflorescence stem, turgescences from vacuole provides support for the cell instead of cellulose in cell walls. Thus, maintaining cell shape during the clearing process is the critical step for clearing seedlings. We modified the PATCSOS protocol by shortening the decolorization/delipidation time and controlling dehydration time strictly to fit the tender nature of seedlings and achieved excellent clearing effect while maintaining cell shape. Hence, the shoot apical meristem (SAM) and trichome can stay in good status after clearing (Fig. 4i, 4j). Eight-day *Col-0* seedlings were cleared with the modified PATCSOS method and loaded onto a glass slide with a central groove. The whole seedling was imaged with a 40×oil objective using 488 nm excitation laser with a step size of 1 μm. The image of the seedling vascular bundle showed that the annular vessel and spiral vessel are the major vessel elements in the vascular bundle (Fig. 4a). An image of part of the shoot apical meristem (SAM) clearly showed cell configuration details of the bud primordium (Fig. 4b). When the laser was focused at the root platform, the root hair could be clearly visualized (Fig. 4c, 4c1), when the laser was focused at the SAM platform, bud primordium could be clearly visualized (Fig. 4d, 4d1), both images were processed by the brightness adjust function of the Fiji software. 3-D imaging of whole seedling showed clear details of all the root hairs, which also proved that small details can be well preserved by processing with PATCSOS (Fig. 4e, 4f, 4g). A movie was made to show more details of the whole seedling (Movie S2). A slice image in the z-direction of the diarch stele in *Arabidopsis* root showed the xylem and phloem arrangement (Fig. 4h). Details of the surfaces of cotyledon and SAM can be clearly visualized, and the structure of the trichomes was also well-preserved (Fig. 4j). The modified PATCSOS method realized whole tissue 3-D imaging of *Arabidopsis* seedling without the need of dissection, which is perfectly suitable for 3-D imaging of small and tender tissues such as seedlings, as such tissues are generally very soft and small in diameter which render them difficult to dissect/slicing.

**Figure 4.**
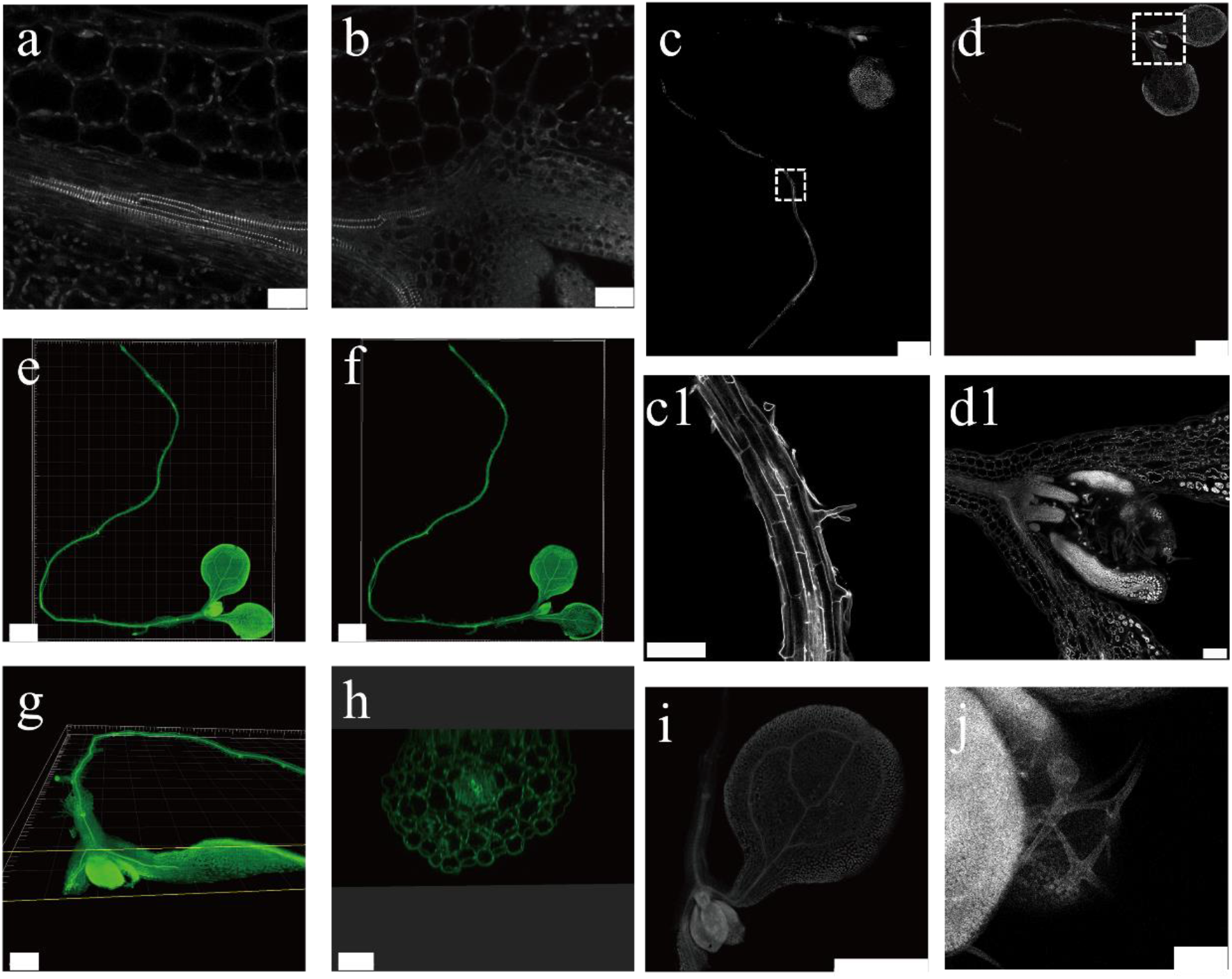
3-D imaging of *Arabidopsis* whole seedling: (a) vascular bundle in hypocotyl; (b) a prat of SAM; (c, d) two images of single light slices of seedling focused on different depths; (c1) root hairs, a magnification of the white dashed box in (c); (d1) bud primordium, a magnification of the white dashed box in (d); (e-g) different views of whole seedling 3-D image; (h) Z direction view of the seedling root; (i) an image of cotyledon; (j) an image of trichome. Bar, 1 μm (a, b); Bar, 1 mm (c, d, e, f, i); bar, 100 μm (c1, d1, j); bar, 500 μm (g); bar, 300 μm (h).

### 3.5 GUS staining after clearing

GUS staining is a widely used technology to reveal gene expression position and expression level. But in some non-cleared tissues, GUS staining may be difficult to observe due to the opaque nature of plant tissues. Thus, the combination of GUS staining and tissue clearing is a perfect way to realize GUS staining from deep-buried, opaque tissue regions. Here, we tested the compatibility of the PATCSOS system with GUS staining. Eight-day seedlings were first stained with conventional GUS staining solution [19]. To adapt to the PATCSOS system, the tissues were decolorized at 30%, 50%, and 70% tB for 10 minutes, respectively, instead of ethanol in the conventional GUS staining protocol. The decolorized seedling can be cleared in less than 12 h by immersing in a standard PATCSOS clearing solution.

*HB29* is a gene we were concerned about at the earlier time. And *DR5* is a famous auxin respond element.[20] So, we chose these two genes to test our protocol. *HB29Pro::GUS* seedling was first GUS stained and then processed with PATCSOS-GUS clearing, the difference in contrast between tissues is more clearly (Fig. 5a, 5c). *DR5::GUS* seedling were also treated with GUS staining and then with PATCSOS-GUS clearing (Fig. 5b, 5d), some edge areas with unclear coloring in traditional GUS staining were now clearly visible. Thus, the PATCSOS-GUS clearing system can maintain GUS staining after clearing and the higher contrast between GUS staining and cleared tissue makes high resolution 3-D imaging available.

**Figure 5.**
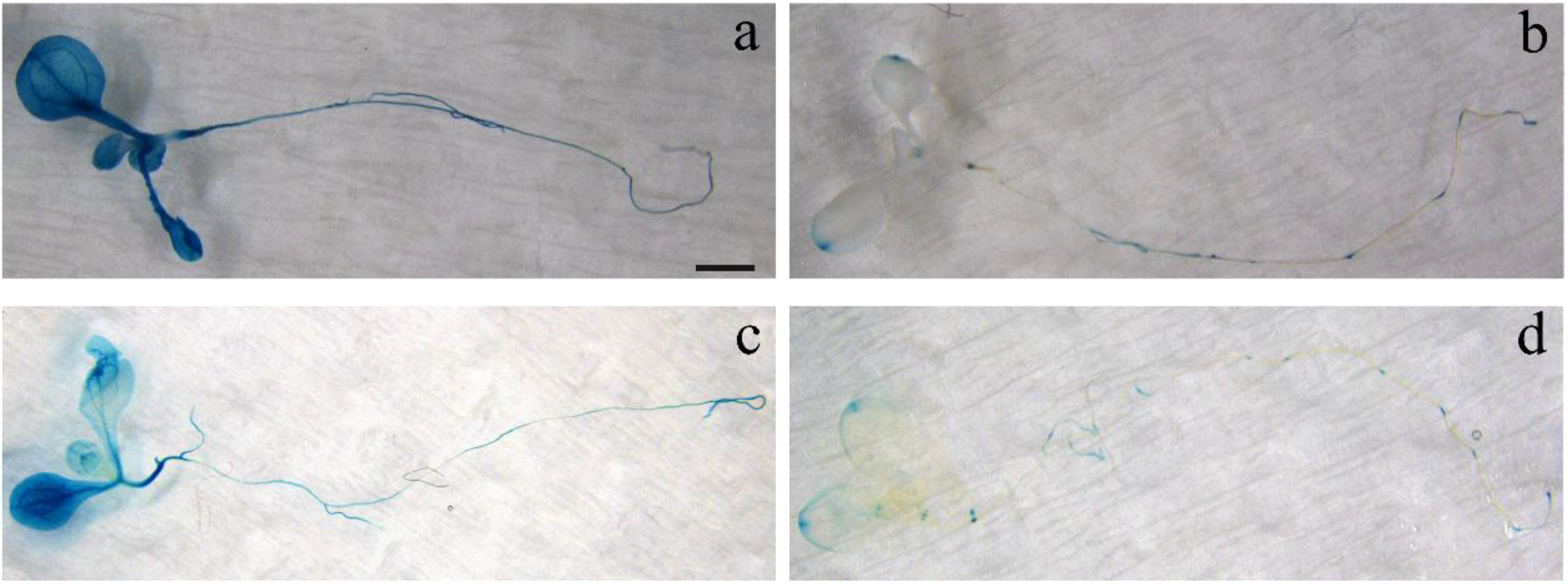
Comparison between traditional GUS staining and PATCSOS-GUS staining: (a) *HB29Pro::GUS* stained by the traditional method; (b) *DR5::GUS* stained by the traditional method; (c) *HB29Pro::GUS* stained by the PATCSOS-GUS method; (d) *DR5::GUS* stained by the PACTSOS-GUS method. Bar, 1 mm.

## 4 Conclusions and Discussion

In this work, we modified the popular mouse tissue clearing method PEGASOS to suit application in plant science research and developed the PATCSOS protocol. The PATCSOS procedure is FPs friendly and can maintain tissue structure of tender *Arabidopsis* seedlings. We also confirmed that PATCSOS is compatible with GUS staining using the modified PATCSOS-GUS protocol.

With the developed method, 3-D images of different plant organs were acquired. A new concept for autofluorescence imaging was proposed, which uses autofluorescence from cell wall components for 3-D imaging, and the autofluorescence signal has generally been considered as noise signal. With autofluorescence, 3-D image from an intact seedling was obtained. By acquiring fluoresce signals from different excitation channels, fluorescence signals from autofluorescence and FPs can be discriminated to reveal “true” signal from the FPs, in this way we imaged the cambium pattern in *Arabidopsis* stipe.

The PATCSOS protocol still has some disadvantages, tB was used for decolorization/delipidation, but, compared with ethanol, the chlorophyll removing efficiency of tB is much lower, but ethanol will quench the FPs fluorescence. So, it is necessary to find a fluorescence friendly and effective chlorophyll removing reagent. And for seeds, which is regarded as the most difficult organ to be cleared, our current protocol still has difficulties to fully clear seeds. Light-sheet microscopy is an ideal technology to image whole mature plant, but we found that the light beam was scattered badly by the vascular tissue. Thus, a better method to control light scattering by the vascular bundle is needed to realize 3-D imaging with light-sheet.

## Supporting information

all supplemental data

## 5 Supplementary Materials

The following are available in supplementary document.

Table S1: Primers used in this study;

Table S2 GUS assay solution (50 ml);

Table S3 X-Gluc mother liquor (250 μL).

Movie S1: 3-D imaging of Silique;

Movie S2: Whole seedling 3-D imaging presentation.

## 6 Author Contributions

Zhang HT. designed the project, performed the experiments and imaging, and wrote the original draft; Li Z., Zhao H., Zhu L. reviewed and edited the article. All authors have read and agreed to the published version of the manuscript.

## 7 Acknowledgments

We thank Beijing Students’ Platform for innovation and entrepreneurship training program for providing apart of financial support. We also thank Yutao Wang, Huan Zhao, Hongjie Xing (China Agricultural University) for help in construction of transgenic plant material, and Youqi Li, Yuling Wang, Manyu Chen, Jiayi Ding (Chinese Institute for Brain Research, Beijing) for the imaging technology assistance.

## 8 Conflicts of Interest

The authors declare no conflict of interest.

## Notes

### Competing Interest Statement

The authors have declared no competing interest.

