## Supplementary material for "Plant Accessible Tissue Clearing Solvent System (PATCSOS) for 3-D Imaging of Whole Plants": all supplemental data

Table S1 Primers used in this study

| Primer name | Sequence (5’→3’) |
| --- | --- |
| ANT Pro for 1300 F | CTTGCATGCCTGCAGGTCGACTATGTTTCCACGTAAAGTTTGGAGG |
| ANT Pro for 1300 R | GAGCTCGGTACCCGGGGATCCGTGTTCTTATTGAGAAAGACGTTTC |
| HB29 Pro for GUS F | TGGCTGCAGGTCGACGGTCATTGACAAGATCGTTAGGAGA |
| HB29 Pro for GUS R | GGTGGACTCCTCTTAAGCTTTTGTTTAGTTCTGTCTTTAT |

Table S2 GUS assay solution (50 ml)

| Reagent | Density | Volume | Company |
| --- | --- | --- | --- |
| NaH_2_PO_4_ | 0.2 M | 7 mL | HUSHI 20040718 |
| Na_2_HPO_4_ | 0.2 M | 18 mL | HUSHI 10020318 |
| K_3_[Fe(CN)_6_] | 0.5 M | 5 μL | HUSHI 10016718 |
| K_4_[Fe(CN)_6_] | 0.5 M | 5 μL | SCRC 10016818 |
| EDTA | 0.5 M | 1 mL | Sigma-Aldrich E5134 |
| TritonX-100 | 10% | 50 μL | Sigma-Aldrich T8787 |
| X-Gluc mother liquor |  | 500 μL |  |
| DdH_2_O |  | 23.44 μL |  |

Table S3 X-Gluc mother liquor (250μL)

| Reagent | Volume or Weight | Company |
| --- | --- | --- |
| N,N-Dimethylformamide（DMF） | 250 μL | XiLONG SCIENTIFIC 12503501 |
| X-Gluc powder | 25 mg | Biotopped X6060 |

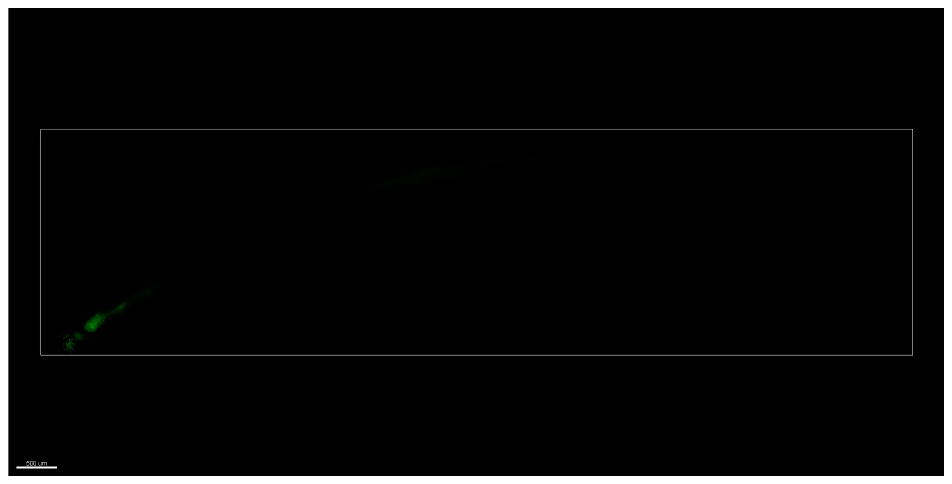

Movie S1 3-D imaging of Silique. Autofluorescence of cellulose was excited using 488-nm argon lasers, and imaged with a 20× objective. Reconstructed 3-D images of an intact silique, slices in both X-Y direction and Z direction were shown. And configuration of seeds in the silique was also shown in this movie.

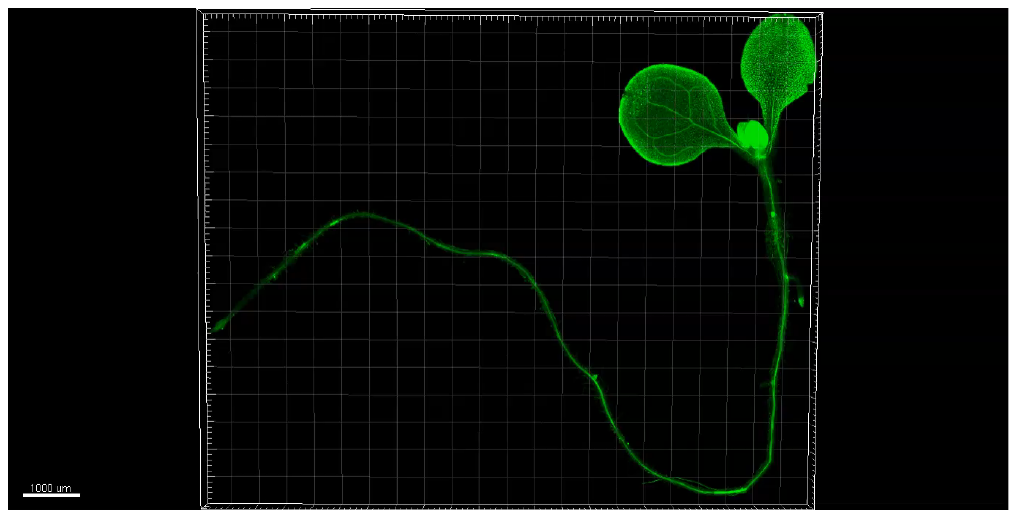

Movie S2 Whole seedling 3-D imaging presentation. Autofluorescence of cellulose was excited using 488-nm argon lasers, and imaged with a 40× objective. Reconstructed 3-D images of whole seedling, slices in both X-Y direction and Z direction were shown. The SAM and root tips were shown especially in this movie.
